# Dopamine-dependent loss aversion during effort-based decision-making

**DOI:** 10.1101/714840

**Authors:** Xiuli Chen, Sarah Voets, Ned Jenkinson, Joseph M. Galea

**Affiliations:** School of Psychology, University of Birmingham, Birmingham, UK, B15 2TT; School of Sport, Exercise and Rehabilitation Sciences, University of Birmingham, Birmingham, UK, B15 2TT

## Abstract

From psychology to economics there has been substantial interest in how costs (e.g., delay, risk) are represented asymmetrically during decision-making when attempting to gain reward or to avoid punishment. For example, in decision-making under risk, individuals show a tendency to prefer to avoid punishment than to acquire the equivalent reward (loss aversion). Although the cost of physical effort has received significant recent attention due to the evaluation of motor costs being crucial in our daily decisions, it remains unclear whether loss aversion exists during effort-based decision-making. On the one hand, loss aversion may be hardwired due to asymmetric evolutionary pressure on losses and gains and therefore exists across decision-making contexts. On the other hand, distinct brain regions are involved with different decision costs, making it questionable whether similar asymmetries exist. Here, we demonstrate that young healthy participants exhibit loss aversion during effort-based decision-making by exerting more physical effort in order to avoid punishment than to gain a same-size reward. Next, we show that medicated Parkinson’s disease (PD) patients show a reduction in loss aversion compared to age-matched controls. Behavioural and computational analysis revealed that people with PD exerted similar physical effort in return for a reward, but were less willing to produce effort in order to avoid punishment. Therefore, loss aversion is present during effort-based decision-making and can be modulated by altered dopaminergic state. This finding could have important implications for our understanding of clinical disorders that show a reduced willingness to exert effort in the pursuit of reward.

**Significance Statement:** Loss aversion – preferring to avoid punishment than to acquire equivalent reward – is an important concept in decision-making under risk. However, little is known about whether loss aversion also exists during decisions where the cost is physical effort. This is surprising given that motor cost shapes human behaviour, and a reduced willingness to exert effort is a characteristic of many clinical disorders. Here, we show that healthy individuals exert more effort to minimise punishment than to maximise reward (loss aversion). We also demonstrate that loss aversion is modulated by altered dopaminergic state by showing that medicated Parkinson’s disease patients exert similar effort to gain reward but less effort to avoid punishment. Therefore, dopamine-dependent loss aversion is crucial for explaining effort-based decision-making.

## Introduction

In daily life we make countless choices that involve a cost-benefit analysis. As a result, there has been substantial interest into how a cost, such as the delay and uncertainty of reward, alters the subjective value an individual associates with the beneficial outcome of a decision (Bautista, Tinbergen, & Kacelnik, 2001; Fehr & Rangel, 2011; Kahneman & Tversky, 1979; Stephens, 2001; Stephens & Krebs, 1986; Stevens, Rosati, Ross, & Hauser, 2005). As many decisions we make requires an evaluation of the associated motor costs (Klein-Flügge, Kennerley, Friston, & Bestmann, 2016), one cost that has received significant recent attention is physical effort (effort-based decision-making). Previous work has investigated the computational, neural and neurochemical mechanisms involved when individuals evaluate rewards that are associated to physical effort (Burke, Brunger, Kahnt, Park, & Tobler, 2013; Hauser, Eldar, & Dolan, 2017; Irma Triasih Kurniawan, Guitart-Masip, & Dolan, 2011; Prévost, Pessiglione, Météreau, Cléry-Melin, & Dreher, 2010), with a diminished willingness to exert effort being a prevalent characteristic of many clinical disorders such as Parkinson’s disease (Baraduc, Thobois, Gan, Broussolle, & Desmurget, 2013; Chong et al., 2015).

With other costs, such as delay and uncertainty, prior work has examined how they are represented differently when attempting to gain reward or avoid punishment. For example, in decision-making under risk, individuals show a tendency to prefer to avoid punishment than to acquire the equivalent reward, a phenomenon called loss aversion (Kahneman & Tversky, 1979; Tversky & Kahneman, 1992). Surprisingly, it remains unclear whether people also exhibit loss aversion during effort-based decision-making. On the one hand, loss aversion may be hardwired due to asymmetric evolutionary pressure on losses and gains (Kahneman & Tversky, 1979; Tom, Fox, Trepel, & Poldrack, 2007; Tversky & Kahneman, 1992b), and thus should be observed in any cost-benefit decision-making context. On the other hand, distinct brain regions are involved in decision-making with different costs (Bailey, Simpson, & Balsam, 2016; Galaro, Celnik, & Chib, 2019; Hauser et al., 2017; Prévost et al., 2010), making it questionable whether similar asymmetries should exist. For example, the critical neural signature of effort-based decision-making is the cingulate cortex, and not the ventromedial prefrontal cortex as typically described for decision-making under risk (Klein-Flügge et al., 2016). Although several studies have attempted to address this question, these either do not directly examine loss aversion (Galaro et al., 2019), do not involve the execution of the effortful action (Nishiyama, 2016) or the cost of effort is confounded with the cost of temporal delay (Porat, Hassin-Baer, Cohen, Markus, & Tomer, 2014).

The neurotransmitter dopamine appears to be crucial for effort-based decision-making. For example, Parkinson’s disease (PD) patients off dopaminergic medication exhibit a reduced willingness to exert effort in the pursuit of reward, with medication restoring this imbalance (Chong et al., 2015; Le Bouc et al., 2016; Skvortsova, Degos, Welter, Vidailhet, & Pessiglione, 2017). Interestingly, during decision-making under risk and reinforcement learning, Parkinson’s disease patients on dopaminergic medication display an enhanced response to reward but a reduced sensitivity to punishment (Collins & Frank, 2014; Frank, 2005; Frank, Seeberger, & O’Reilly, 2004). Although this suggests that dopamine availability might shape loss aversion across contexts (Clark & Dagher, 2014; Timmer, Sescousse, Esselink, Piray, & Cools, 2017), and in particular that medicated PD patients should show reduced loss aversion, the role of dopamine during effort-based decision-making within a reward or punishment context has not been directly investigated.

In this paper, we demonstrate that young healthy participants exhibit loss aversion during effort-based decision-making; individuals were willing to exert more physical effort in order to minimise punishment than maximise reward. In addition, behavioural and computational analysis revealed that medicated Parkinson’s disease patients showed a reduction in loss aversion compared to age-matched controls. Specifically, although patients exerted similar physical effort in return for reward, they were less willing to produce effort to avoid punishment. Therefore, loss aversion is present during effort-based decision-making and this asymmetry is modulated by dopaminergic state.

## Materials and Methods

### Participants

#### Ethics statement

The study was approved by Ethical Review Committee of the University of Birmingham, UK, and was in accordance with the Declaration of Helsinki. Written informed consent was obtained from all participants.

#### Young healthy participants

Twenty-two young healthy participants (age: 23.1 ± 4.56; 16 females) were recruited via online advertising and received monetary compensation upon completion of the study. They were naïve to the task, had normal/corrected vision, and reported to have no history of any neurological condition.

#### Parkinson’s disease patients (PD) and healthy age-matched controls (HC)

Eighteen PD patients were recruited from a local participant pool through Parkinson’s UK. They were on their normal schedule of medication during testing (levodopa-containing compound: n=7, dopamine agonists (including pramipexole, ropinirole): n=6, or combination of both: n=5). Clinical severity was accessed with the Unified Parkinson’s Disease Rating Scale (UPDRS, Table 1) (Fahn & Elton, 1987). Twenty HC were also recruited via a local participant pool. Both groups received monetary compensation upon completion of the study. All patients/participants had a Mini-Mental Status Exam (Folstein, Folstein, & McHugh, 1975) score greater than 25 (Table 1). Table 1 summarises the demographics of the patients and age-matched controls. For additional motivation, all patients/participants were offered a chance to enter a lottery to win an extra £100 if their performance (final score) was among the top 5 (one lottery per group).

**Table 1:**
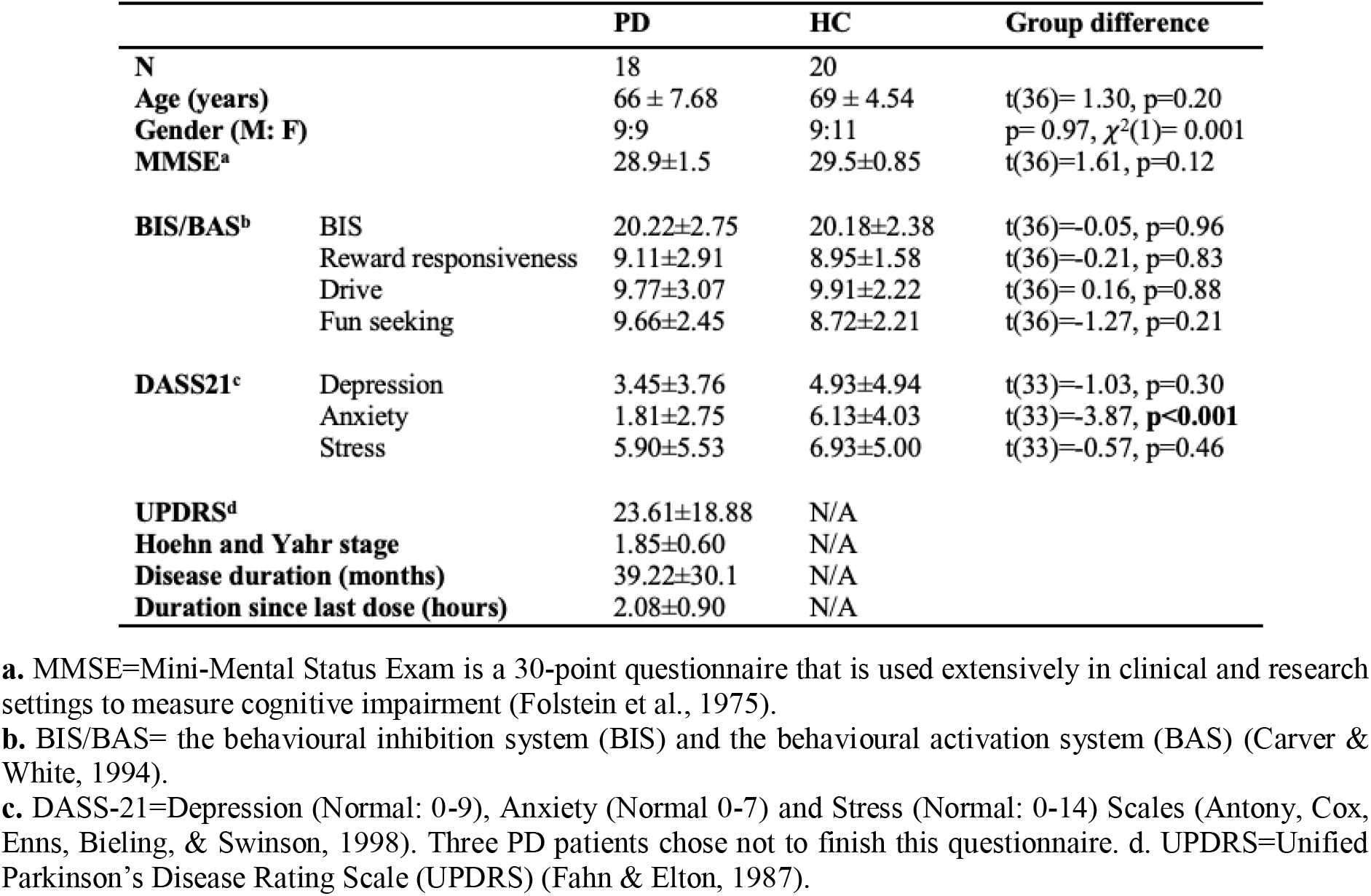
Demographics for PD and HC groups (means ± SD)

### Experimental design

#### Experiment set up

Participants were seated in front of a computer (Figure 1A) running a task implemented in Psychtoolbox (http://Psychtoolbox.org) and Matlab (MathWorks, USA). Two custom-built vertical handles were positioned on a desk in front of the participants, each of which housed a force transducer with sample rate of 200 hertz (https://www.ati-ia.com). The force produced on each handle enabled participants to independently control two cursors on the computer screen (Figure 1A). During the main experiment, one handle was assigned as the decision-making handle; participants grasped this handle with their hand and produced a left or right directed force in order to move the decision cursor into the appropriate option box to indicate their choice. The other handle was designated as the force execution handle; participants rested their index finger next to the bottom of the handle and produced a force by pressing their index finger inward on the handle (i.e., push left for the right index finger, push right for the left index finger).

**Figure 1:**
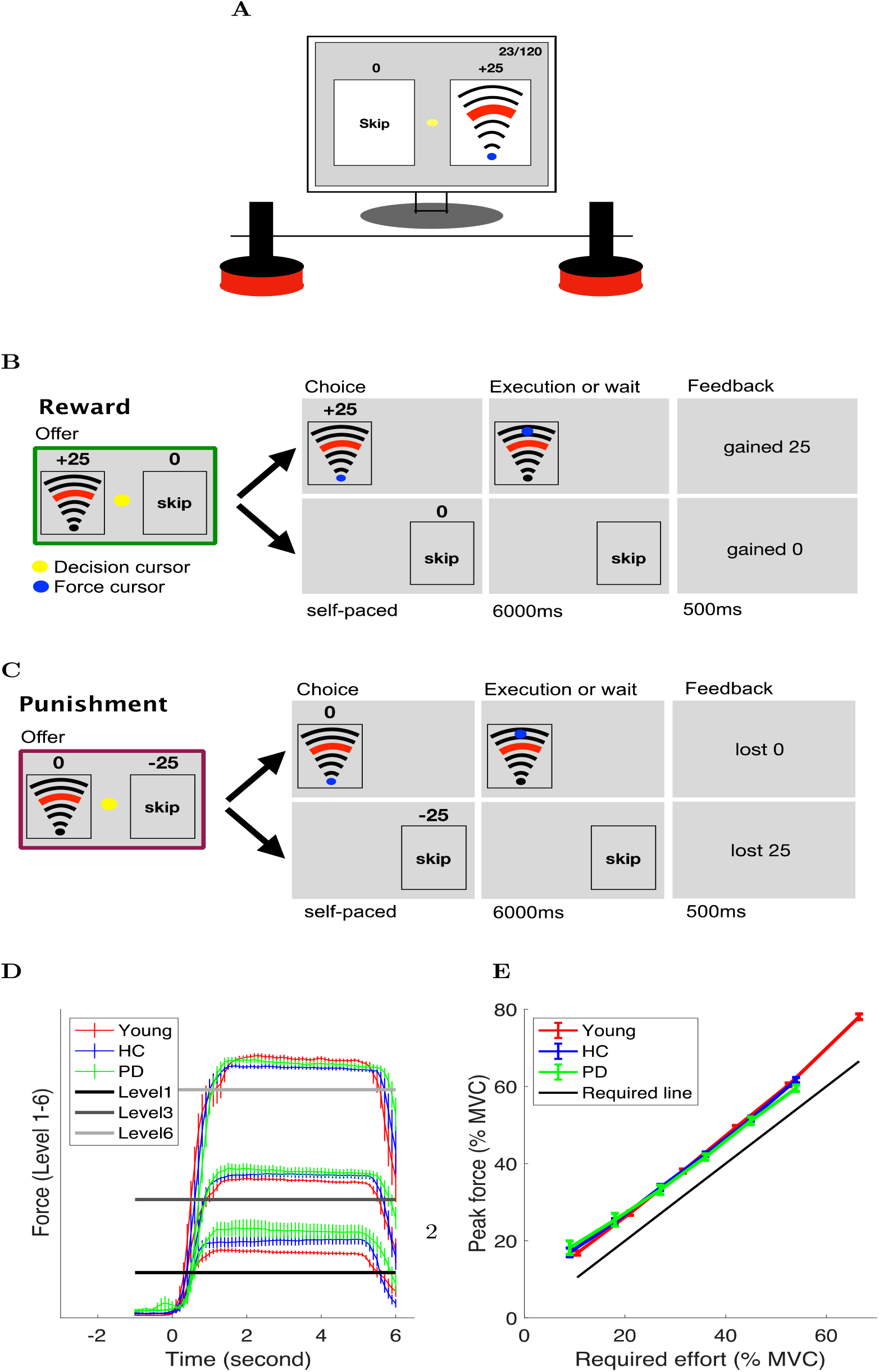
Experimental setup. **(A)** Experimental equipment. **(B-C)** Typical reward (B) and punishment (C) trial. **(D)** Average force trace across participants on levels 1, 3 and 6. 0 second (x-axis) is the moment at which the participants indicated their choice and they were allowed to start exerting the force. **(E)** Young participants (red), PD patients (green) and healthy age-matched controls (blue) all modulated their force appropriately. The solid black line indicates the minimum required force.

#### Procedure

Before the main effort-based decision-making task, participants were asked to produce a maximal voluntary contraction (MVC) of their first dorsal interosseous (FDI) muscle (isometric contraction of the index finger against the handle) for 3 seconds. This was repeated 3 times and the average maximum force was taken as their MVC. For the young healthy participants, the index finger of the dominant hand was chosen to produce the force. For PD patients, the index finger of the most affected side was chosen to produce the force (dominant hand n=11, non-dominant hand n=7). For the healthy age-matched controls, we chose a similar ratio of dominant hand and non-dominant hand as their force producing hand (dominant hand n=12, non-dominant hand n=8). Following the MVC, participants had 12 trials to practise the 6 force levels that were used in the main decision-making task (see *Effort-based decision-making task* section for details). The force levels were shown to participants as a set of arcs (Figure 1A).

The effort-based decision-making task consisted of 2 blocks (reward and punishment), the order of which was counter-balanced across participants. Each block consisted of 10 repetitions of each of the 6 force levels, with a total of 60 trials in each block (15 repetitions for the young age group, 90 trials in each block). Following the effort-based decision-making task, participants were asked to produce 3 consecutive 3-second MVCs. They were instructed that this had to be within 90% of the MVC they produced at the beginning of the experiment. Importantly, participants were made aware of this requirement at the beginning of the study (after the MVC and before the main decision-making task). This was intended to ensure that they cared about not becoming over fatigued by always choosing the effortful (high reward, low punishment) choice throughout. Therefore, all participants were encouraged to accumulate as many points as possible (and lose as few points as possible), whilst avoiding unnecessary effort.

#### Effort-based decision-making task

The task was adapted from classic effort-based decision-making paradigms (Bonnelle, Manohar, Behrens, & Husain, 2016; Bonnelle et al., 2015; T. T.J. Chong, Bonnelle, & Husain, 2016; Le Heron et al., 2018; Skvortsova et al., 2017). There were two trial types: reward and punishment (Figure 1B,C) and the task consisted of one block of each. On a reward trial (Figure 1B), participants chose between executing a certain force level in return for reward (gaining points) and skipping the trial in return for 0 points. On a punishment trial (Figure 1C), participants chose between executing a certain force level in return for 0 points and skipping the trial in return for being punished (losing points).

On each trial, participants were presented with a combination of points and a force level, which was a percentage of their MVC (offer phase). For the young group, the force was 1 of 6 levels: 11, 21, 32, 42 53, 67% of MVC. For both the older age groups (PD and HC), these six levels were: 9, 18, 27, 36, 45, 54% of MVC. The force levels used for the older age groups were lower because a pilot study revealed they fatigued significantly faster than younger participants. At the beginning of each block (reward, punishment), these six force levels were paired with [5 10 15 20 25 30] points respectively. The initial pairings were selected based on pilot experiments. Unbeknown to participants, the points associated with each force level were then adjusted on a trial-by-trial basis using an adaptive staircase algorithm (Parameter Estimation by Sequential Testing; Taylor & Creelman, 2005). Specifically, the points offered were increased or decreased using an initial step size of 8, depending on whether participants rejected (skipped) or accepted the opportunity to execute the force in order to receive (or avoid losing) those points. The step-size was doubled if participants rejected or accepted the offer (a combination of force and points) 3 times in a row, and the step-size was halved if participants reversed their decision on the force level, i.e., an acceptance followed by a rejection on a force level or vice-versa (Taylor & Creelman, 2005). As the staircase procedure was performed independently for each of the six force levels, it allowed us to determine the point of subjective indifference at which participants assigned equal value to acceptance and rejection for each force level. The trial order was randomized across the 6 force levels. Importantly, the points and force combinations offered in the reward and punishment conditions were under the same adaptive procedure as described above, the only difference being whether the points were framed as rewards or punishments (Figure 1 B,C; Tversky & Kahneman, 1981).

Following the offer phase, participants indicated their choice by exerting a force on the decision handle which moved the yellow decision cursor (Figure 1A) from the middle of the screen into one of the option boxes (execute force or skip trial). As soon as participants indicated their choice, the unchosen option disappeared. If the force option was chosen, participants were required to execute the force on the handle with this being represented by the blue force cursor moving from the start position towards a target line, and staying above the target line for 4 seconds at which point they heard a cash register sound ‘ka-ching’ from the headphone. If they failed to exert the required force, the trial was repeated. The trial was always terminated 6.5 seconds after their choice. This meant that participants had to wait for 6 seconds if they chose to skip the trial, or they had to produce the required force within 6 seconds. We carefully controlled the time for force execution and skip decisions to be identical so that there was no confound between delay and effort discounting as in previous studies (Doyle, 2010; Loewenstein, Frederick, & O’donoghue, 2002).

#### Data and statistical analysis

Data were analysed with Matlab using custom scripts. The data and codes are available at https://osf.io/hw4rk/. Our first question was to ask if young healthy participants expressed loss aversion during effort-based decision-making, i.e., a preference to exert more physical effort in order to minimise punishment than maximise reward. For each of the six force levels, we estimated the points at which the probability of accepting the force option was 50% (effort indifference point). Specifically, for each force level, a logistic function 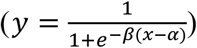 was fitted to the points offered and the binary choices made by participants (Figure 2). As shown in Figure 2, the effort indifference point was then defined as the reward magnitude (x-axis) at which the sigmoid crossed y = 0.5.

**Figure 2:**
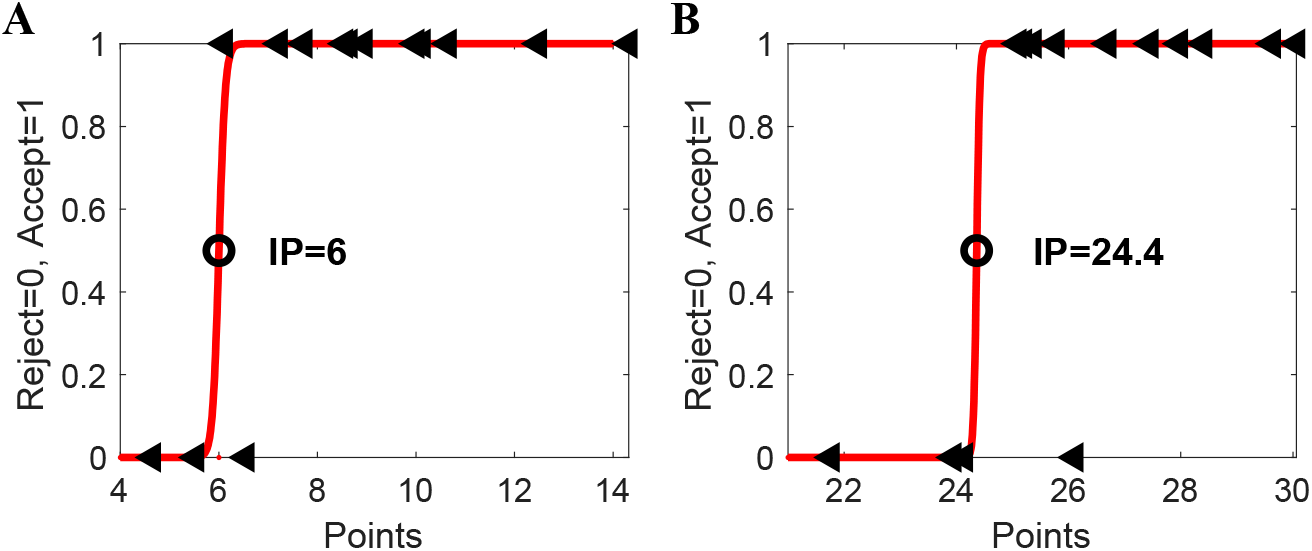
Procedure for determining effort indifference point. Exemplary choices and fits are shown for one participant and two effort levels (**A:** level 2; **B:** level 6) in the reward condition. A sigmoid function (red line) was fitted separately to the choices (arrow) generated at each effort level (y axis: 0 = reject force, 1 = accept force), given the points (reward or punishment) offered for this force level (x-axis). The point of subjective indifference point (IP, circle) was defined as the magnitude at which the sigmoid crossed y = 0.5.

An average effort indifference point (across six force levels) was then calculated for each participant in the reward and punishment conditions (referred to as reward IP and punishment IP respectively), indicating an individual’s tendency to produce force in each condition. Each participant’s loss aversion index was then defined as a ratio between reward IP and punishment IP. A loss aversion index that was larger than 1 indicated loss aversion. Due to non-normalities in the data, a Wilcoxon Signed-ranks test (*signrank* function in Matlab) was used to test if the loss aversion index for young healthy participants was significantly greater than 1. To assess effort-based loss aversion in PD patients and HC, we compared their loss aversion index using non-parametric independent samples Mann-Whitney U-tests (*ranksum* function in Matlab). To examine the loss aversion differences in more detail, a two-way mixed ANOVA compared the average effort indifference point across group (PD vs HC) and condition (reward vs. punishment). In order to address non-linearity and heteroscedasticity (unequal variance), the effort IP was log-transformed.

#### Computational modelling of choice

As with other forms of decision-making, choices during effort-based decision-making depend on cost-benefit analyses. For instance, the subjective value of reward decreases as the physical effort associated with it becomes progressively more demanding (Botvinick, Huffstetler, & McGuire, 2009). This relationship between reward and effort is often captured by effort discounting functions (Białaszek, Marcowski, & Ostaszewski, 2017; Botvinick et al., 2009; Hartmann, Hager, Tobler, & Kaiser, 2013; Klein-Flügge, Kennerley, Saraiva, Penny, & Bestmann, 2015; Prévost et al., 2010). One of the key parameters in an effort discounting function is the discounting parameter, which denotes the steepness of how effort discounts reward. That is, it represents the willingness to invest effort for a beneficial outcome. Therefore, in our effort-based decision-making task, differences in choice behaviours between groups or across reward and punishment conditions could potentially manifest as changes in the discounting parameter of an effort discounting function. To test this, we fitted participant responses using linear, parabolic and hyperbolic effort discounting functions, which are often used to capture effort discounting (Białaszek et al., 2017; Hartmann et al., 2013; Klein-Flügge et al., 2015; McGuigan et al., 2019). The shape of these functions reflects how increasing costs (i.e., effort) discounts the associated benefits (i.e., gaining rewards, avoiding punishments).

The linear model is described by: *SV = A — lE* (McGuigan et al., 2019), where *SV* denotes the subjective value of a reward/punishment *A*. The parameter, *E,* denotes the effort involved in order to gain a reward or to avoid a punishment, which was the percentages of each individual maximum force (MVC). The parameter, *l*, is the steepness of the discounting parameter, which can be interpreted as the unwillingness to exert effort. A higher value of *l* represents less willingness by an individual to expend effort for the given outcomes. The parabolic model is described by: *SV = A — lE^2^* (Hartmann et al., 2013). This function implies that additional effort devalues a reward to a greater extent if existing effort is high rather than low. The hyperbolic model is described by: 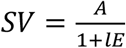 (Mazur, 1987). This function implies that if additional effort is introduced to existing effort, it devalues reward more if the existing effort is low rather than high.

The model space included all possible combinations of linear, parabolic and hyperbolic effort discounting functions in each of the two conditions performed by PD and HC groups. To compare the models, we utilised Bayesian Information Criterion (BIC) (Schwarz, 1978). Specifically, for each model, the BIC summed over all participants were compared (the lower the value, the better the model fit) (Rigoux, Stephan, Friston, & Daunizeau, 2014; Stephan, Penny, Daunizeau, Moran, & Friston, 2009). Such aggregation of BIC across participants corresponds to fixed-effect analyses (Stephan et al., 2009). To account for the random-effect analysis in which models are treated as a random variable that can differ between participants (Stephan et al., 2009), we also conducted Friedman’s test on individual BIC to compare the model fits (non-parametric repeated-measures ANOVA).

## Results

### Evidence for loss aversion in young healthy participants

Our first question was to ask if young healthy participants expressed loss aversion during effort-based decision-making. To examine this, we first assessed how the effort indifference point (Figure 2) was affected by force level in the reward and punishment conditions. As expected, the effort indifference point became progressively larger as the force level became more demanding, indicating a sensitivity to effort across reward and punishment conditions (Figure 3A). For each participant, an average effort indifference point was obtained across force levels for the reward (reward IP) and punishment (punishment IP) conditions, with the loss aversion index being defined as a ratio between these values (>1 = loss aversion; Figure 3B). As the loss aversion index was significantly greater than 1 (z=3.65, p<0.001, median=1.37, Figure 3B), it suggests that loss aversion was clearly evident in young healthy participants during effort-based decision-making.

**Figure 3:**
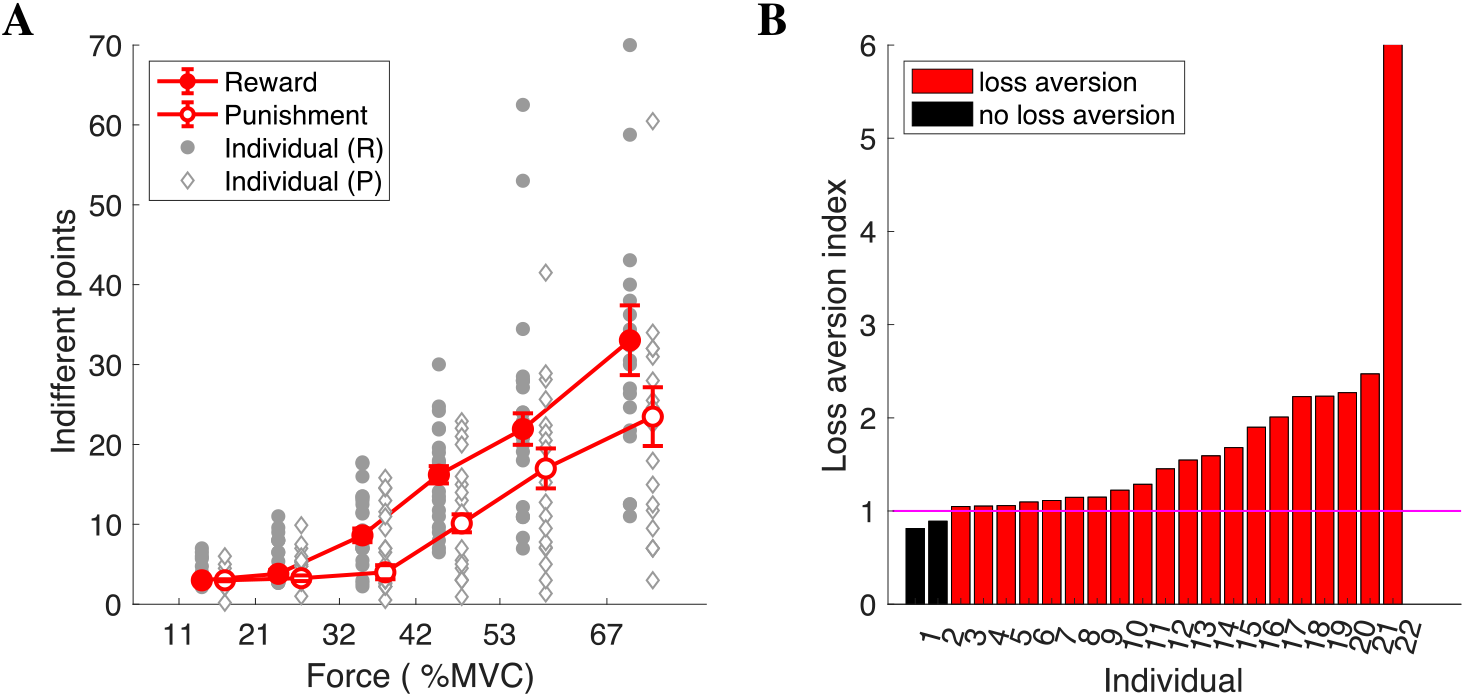
Loss aversion in young healthy participants. **(A)** Effort indifferent points in reward (solid circles) and punishment (open diamonds). For each force level (x-axis), we estimated a score at which the probability of choosing to produce the force was 50% (effort IP, y-axis). Given a particular force level, a higher indifference point indicated less willingness to produce the force. Error-bars represent SEM across participants. Grey circles/diamonds indicate individual data points. **(B)** Loss aversion index for each individual. Loss aversion is reflected by participants being more willing to produce a force to avoid losses than receive same-sized gains (higher reward IP than punishment IP given a force level). Loss aversion was therefore quantified as a ratio between the reward IP and the punishment IP (loss aversion index; y-axis). A value greater than 1 indicates loss aversion.

### Reduced loss aversion in PD patients compared to HC

Similar to the young healthy participants, the effort indifference point for both the HC (Figure 4A) and PD (Figure 4B) groups increased progressively as the force level became more demanding, suggesting sensitivity to effort across reward and punishment conditions. In addition, as the loss aversion index was significantly greater than 1 for both HC (z=3.80, p<0.001, median=2.04, Figure 4C) and PD (z=3.37, p<0.001, median=1.27, Figure 4D), it indicates that loss version was present in both groups. Importantly, PD patients displayed significantly less loss aversion than the HC group (z=2.23, p=0.025, Figure 4E), with this being a result of medicated PD patients appearing less sensitive to punishment (Figure 4F). This was confirmed by a two-way mixed ANOVA which revealed a significant interaction between Group (HC vs PD) and Condition (reward vs punishment) (F[1,36]= 5.22, p=0.028) for the average indifference point. Specifically, Bonferroni-corrected independent t-tests revealed the PD and HC groups had a similar reward IP (p=0.13, Figure 4F), but the PD group displayed a higher punishment IP (p=0.007, Figure 4F).

**Figure 4:**
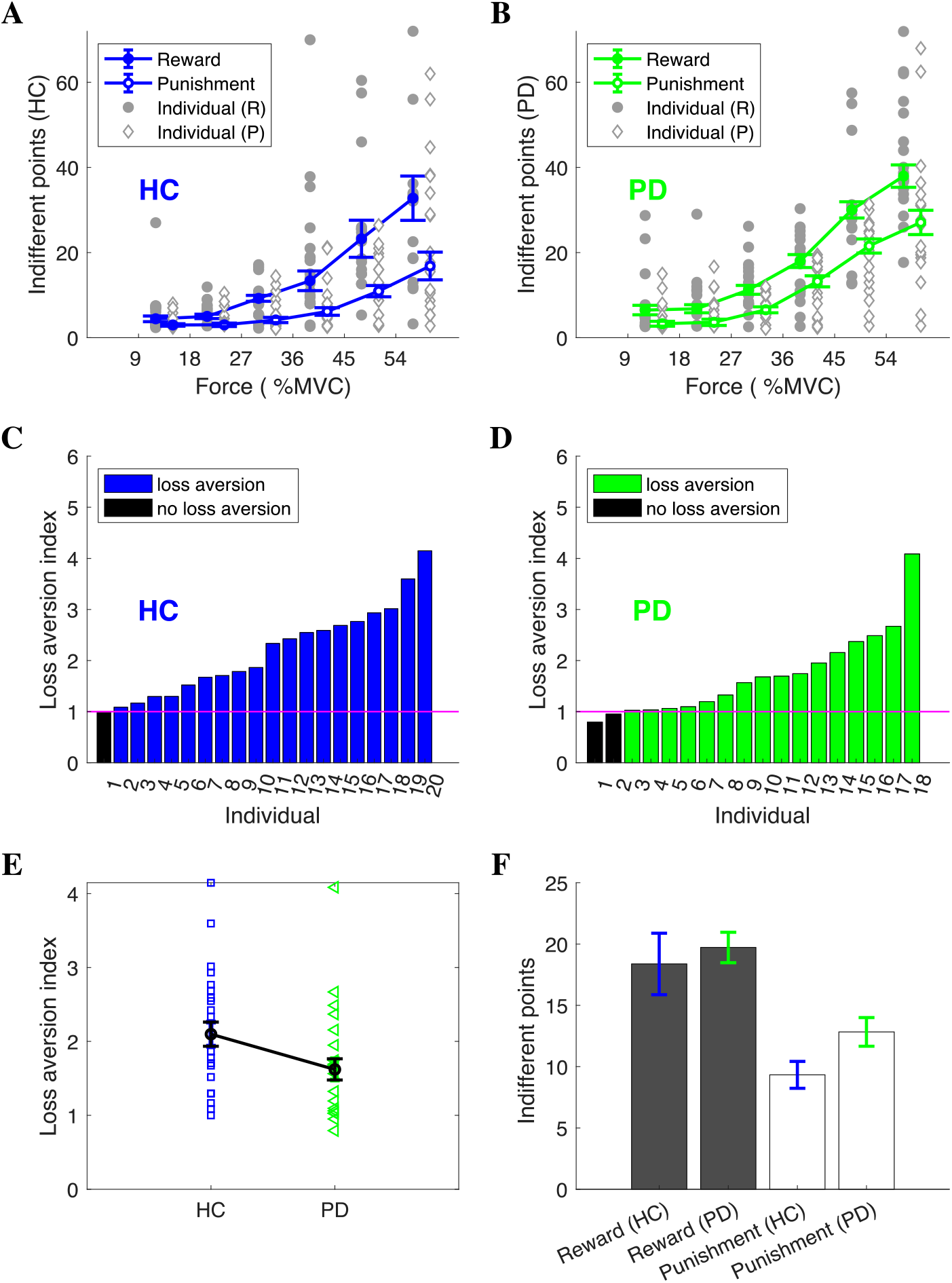
Loss aversion in HC and PD groups. **(A-B)** Effort indifference point in reward (solid circle) and punishment (open diamond) conditions for the HC (**A**) and PD (**B**) groups. For each force level (x-axis), we estimated a score at which the probability of choosing to produce the force was 50% (effort indifference point, y-axis). Given a particular force level, a higher indifference point indicated less willingness to produce the force. Error-bars represent SEM across participants. Grey indicates individual data points. **(C-D)** Loss aversion across participants for the HC (**C**) and PD (**D**) groups. Loss aversion is reflected by participants being more willing to produce a force to avoid losses than receive similar gains. Therefore, the loss aversion index was measured as a ratio between the reward IP and the punishment IP (y-axis). A value greater than 1 indicates loss aversion. **(E)** Loss aversion index. Error-bars represent SEM across participants. **(F)** Reward IP and punishment IP across groups. Steeper effort discounting for PD patients in punishment, but not reward.

The effort discounting parameter represents the steepness of how effort discounts a beneficial outcome, indicating a tendency to expend effort in pursuit of reward or avoid punishment. Therefore, differences in choice behaviours between groups or across conditions could potentially manifest as changes in the gradient of an effort discounting function. To test this, we applied computational models of choice behaviours to estimate the subjective value of each offer to each individual. We fitted participant choices to three typical discounting functions. The model space included all possible combinations of linear, parabolic and hyperbolic effort discounting functions in each of the two conditions performed by PD and HC groups. We found that a parabolic effort discounting function provided the best fit for both the PD and HC groups across the reward and punishment conditions (Table 2). Specifically, the summed Bayesian Information Criterion (BIC) was lowest for the parabolic function (the lower the value, the better the model fit) (Rigoux et al., 2014; Stephan et al., 2009) across groups (HC, PD) and conditions (reward, punishment) (Table 2). To investigate this at a subject-level, a Friedman’s test on individual BIC was performed. In general, similar results were observed with the parabolic function consistently being associated with significantly lower BIC across groups and conditions (Table 2). To reinforce these results, R^2^ was found to be greater for the parabolic function across all groups and conditions (Table 2). Therefore, there appeared to be no difference in the fundamental pattern of effort discounting between groups or conditions.

**Table 2:** Model comparison. The parabolic effort discounting function provided the best fit for choices of both the PD and HC groups across the reward and punishment conditions. Summed BIC, Friedman’s test and R^2^ (mean ± SD) are provided for each group (HC, PD) and condition (reward, punishment).Specifically, for each model, the Bayesian Information Criterion (BIC) summed over all participants were compared (the lower the value, the better the model fit) (Rigoux et al., 2014; Stephan et al., 2009).

|  |  |  | Linear | Parabolic | Hyperbolic | Friedman test |
| --- | --- | --- | --- | --- | --- | --- |
| PD | Reward | BIC | 1529 | 1436 | 1541 |  |
| | | Mean rank | 2.28 | 1.39 | 2.33 | $\chi^2=10.1$; $p=0.006$ |
| | | $R^2$ | $0.77\pm0.21$ | $0.84\pm0.19$ | $0.77\pm0.20$ | |
|  | Punishment | BIC | 1374 | 1319 | 1430 |  |
| | | Mean rank | 2.22 | 1.78 | 2.00 | $\chi^2=1.78$; $p=0.411$ |
| | | $R^2$ | $0.70\pm0.27$ | $0.76\pm0.26$ | $0.71\pm0.24$ | |
| HC | Reward | BIC | 1607 | 1530 | 1611 |  |
| | | Mean rank | 2.35 | 1.40 | 2.25 | $\chi^2=10.9$; $p=0.004$ |
| | | $R^2$ | $0.73\pm0.19$ | $0.79\pm0.21$ | $0.70\pm0.20$ | |
|  | Punishment | BIC | 1287 | 1257 | 1462 |  |
| | | Mean rank | 1.95 | 1.45 | 2.60 | $\chi^2=13.3$; $p=0.001$ |
| | | $R^2$ | $0.54\pm0.26$ | $0.64\pm0.27$ | $0.56\pm0.25$ | |

Using the winning model (parabolic function), we compared parameters across the PD and HC groups. In the reward condition, the effort discounting parameter was found to be similar between the HC and PD groups, suggesting medicated PD patients were equally as motivated to exert effort in return for reward (Figure 5A,B). However, in the punishment condition, the PD groups had an increased effort discounting parameter suggesting they were less willing to exert effort in order to avoid punishment (Figure 5A,C). This was confirmed by a two-way mixed ANOVA that showed a significant interaction between group (HC vs PD) and condition (reward vs punishment) (F(1,37)=6.26, p=0.017). Bonferroni-corrected independent t-tests revealed that while the discounting parameter (*l*) was similar between PD and HC (p=0.342) for reward, it was significantly higher for the PD group in the punishment condition (p=0.032, Figure 5A).

**Figure 5:**
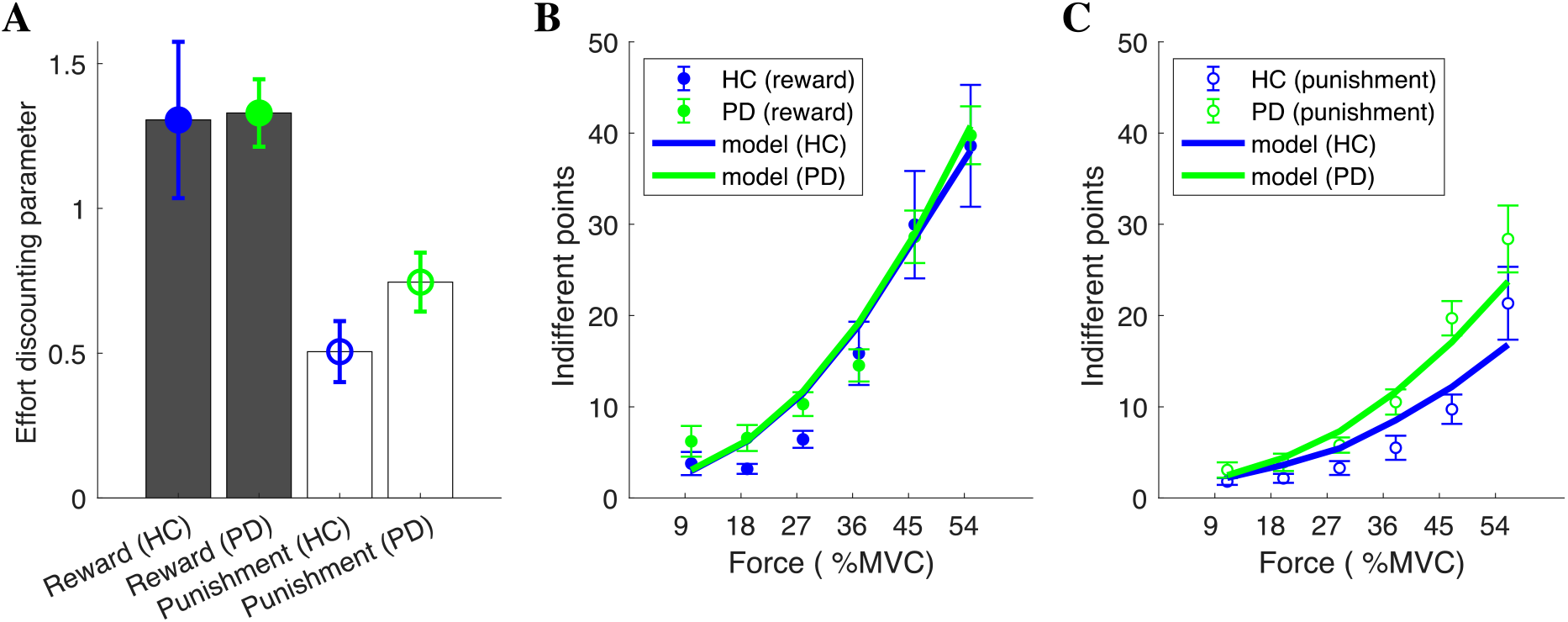
Parabolic (winning model) discounting parameter (l) for the HC and PD groups. **(A)** Effort discounting parameter (*l*) for the HC and PD groups in the reward and punishment conditions. **(B, C)** Parabolic model predictions for the effort indifference point across force options in the reward (**B**) and punishment (**C**) conditions. The model predictions were calculated by estimating a score for which the probability of the model choosing the force option was 50%.

## Discussion

In summary, we have shown that loss aversion is consistently present during effort-based decision-making in young healthy participants and both people with Parkinson’s disease and healthy older adults. Although loss aversion is widely regarded as one of the most robust and ubiquitous findings in economic decision-making (Kahneman & Tversky, 1979; Tversky & Kahneman, 1992), the surprisingly few studies that have directly examined loss aversion during physical effort-based decision-making have found it to not exist. For instance, Porat et al., (2014) showed that while half of young healthy participants were willing to expend greater effort to avoid punishment than to gain an equivalent reward, the other half showed the opposite preference. In addition, Nishiyama, (2016) found a similarly large degree of variability across participants in preference for maximising gains or minimising losses during an effort-based decision-making task. Therefore, while both studies found differences between gain and loss at an individual level, they did not find loss aversion during effort-based decision-making at a group level. However, we believe that there are several issues with the previous studies which may restrict their capacity to directly examine loss aversion during effort-based decision making. First, in Porat et al., (2014) gaining reward or avoiding punishment required the participant to execute additional key presses. As a result, to obtain more reward (or avoid more punishment) the participants had to produce more effort and also had to wait longer. Therefore, the additional effort cost was always confounded with a temporal delay cost. It is worth noting that the temporal discount for losses are generally less steep than that for gains (Estle, Green, Myerson, & Holt, 2006). Importantly, this confound was carefully eliminated in our paradigm as all trials, including the skip option trials, had identical durations. Second, in Nishiyama, (2016), participants were tasked with making a series of choices of whether to engage in an effortful task (to obtain reward or to avoid punishment) via a questionnaire. That is, participants did not actually have to perform an effortful task. The absence of loss aversion could be a result of participants being less sensitive to the imaginary effort involved in a questionnaire. This possibility is supported by our results in which loss aversion is more clearly expressed at higher effort levels.

The second key finding of the present study was that people with Parkinson’s Disease (PD) on medication showed a reduction in loss aversion compared to age-matched healthy controls. Importantly, we found that this reduction in loss aversion was due to people with PD investing similar physical effort in return for a reward but being less willing to produce effort to avoid punishment compared to aged-matched healthy controls. Although previous studies have already demonstrated that medicated PD patients are equally as motivated to exert effort in return for reward as age-matched controls (Chong et al., 2015; Le Heron et al., 2018; McGuigan et al., 2019), this is the first study to reveal that medicated PD patients exhibit reduced loss aversion during effort-based decision making as a result of a specific reduction in their willingness to produce effort to avoid punishment.

To understand this reduced loss aversion in medicated PD patients, one key question is whether it is due to an altered sensitivity to the cost of effort, an altered sensitivity to the action outcomes or a combination of both. It has been repeatedly shown that PD patients exhibit reduced willingness to expend effort in return of a reward, and dopaminergic medication is able to ameliorate this deficit (Chong et al., 2015, Le Heron et al., 2016, Skvortsova et al., 2017). Many earlier studies have also shown that manipulating dopamine can shift the effort/reward trade-off in healthy participants and animals (Bardgett, Depenbrock, Downs, Points, & Green, 2009; Chong et al., 2015; Floresco, Tse, & Ghods-Sharifi, 2008; J. D. Salamone, Correa, Farrar, & Mingote, 2007). However, despite dopamine being clearly central to effort-based decision-making, its precise role is unclear. This uncertainty is because an increased sensitivity to reward or a decreased sensitivity to effort could both explain a similar shift in preference, and the aforementioned studies have been unable to detangle these two possibilities. Recent work has appeared to come to a consensus that dopamine activity have a limited influence on effort cost evaluation during effort-based decision making, whilst there is strong association between dopamine and the action outcome (Le Bouc et al., 2016; Skvortsova et al., 2017; Walton & Bouret, 2019). That is, even if dopamine seems to promote energy expenditure, it only does so as a function of the upcoming reward and not as a function of the upcoming energy cost itself (Le Bouc et al., 2016; Skvortsova et al., 2017). For example, Le Bouc et al., (2016) showed that the bias toward large reward/high effort options under dopaminergic medication is best captured by a model that indicates dopamine increasing sensitivity to action outcomes and not by it decreasing sensitivity to effort costs. Our data also showed that PD patients did not show a generalised reduction in their willingness to engage in effort across the reward and punishment conditions. Therefore, it seems plausible that the altered choice behaviour in the PD group was not due to an increase in effort sensitivity but was predominantly driven by an altered sensitivity to the expected action outcomes.

Consequently, the reduced loss aversion in medicated PD patients during our effort-based decision-making task could be due to dopamine availability modulating an individual’s sensitivity to reward and punishment-based action outcomes. In the domain of reinforcement learning, dopamine manipulation studies in healthy participants and PD patients have revealed that the balance between learning from reward and punishment is strongly modulated by dopamine availability (Cools, Altamirano, & D’Esposito, 2006; Frank, 2005; Frank et al., 2004). Specifically, increases in dopamine enhance reward-based learning while impairing punishment-based learning, and decreases in dopamine enhance punishment-based learning while impairing reward-based learning. Relatively recently, studies have extended these effects of dopamine from reinforcement learning to decision-making under risk, showing that dopamine modulation can influence an individual’s sensitivity to positive versus negative action outcomes in ways very similar to the dopaminergic effects on learning (Collins & Frank, 2014; Shiner et al., 2012; Smittenaar et al., 2012). This suggests that dopamine availability might shape loss aversion across contexts by changing an individual’s sensitivity to reward-and punishment-based action outcomes (Clark & Dagher, 2014; Timmer et al., 2017). Therefore, in the context of the current study, medicated PD patients exhibited reduced loss aversion because they had a normal sensitivity to reward-based action outcomes but a reduced sensitivity to punishment-based action outcomes. However, demonstrating that the dissociable influence of dopamine availability on reward- and punishment-based action outcome sensitivity is independent of context would require further investigation. Specifically, PD patients would need to show similar changes in loss aversion across decision-making under risk and effort-based decision-making tasks, and this would need to be modulated by medication state.

To counter this argument though, previous work has shown that the benefit/cost analysis with different decision costs (e.g., effort, risk and delay) involve separable brain regions. For example, the critical neural signature of effort-based decision-making has been reported in the cingulate cortex, and not the ventromedial prefrontal cortex as typically described for decision-making under risk (Klein-Flügge et al., 2016). Alternatively, therefore, the current results could also be explained by effort being represented by separable dopamine-dependent brain regions when associated with reward or punishment-based action outcomes, rather than dopamine specifically influencing action outcome sensitivity. For instance, multiple studies have shown that when associated with reward effort is evaluated by dopamine-dependent brain regions such as the cingulate cortex, putamen and supplementary motor area (SMA) (Bonnelle et al., 2016; Hauser et al., 2017; Klein-Flügge et al., 2016). Although no study has investigated how effort is represented when associated with punishment, both the dorsal anterior cingulate cortex and anterior insula are involved in effort (Croxson, Walton, O’Reilly, Behrens, & Rushworth, 2009; I. T. Kurniawan, Guitart-Masip, Dayan, & Dolan, 2013; Rudebeck et al., 2006; Walton, Bannerman, & Rushworth, 2002), and independently punishment processing (Nitschke et al., 2006; Palminteri, Khamassi, Joffily, & Coricelli, 2015). Therefore, there is at least a suggestion, that similar brain regions are involved in the processing of effort and punishment which are independent of the reward system.

In conclusion, loss aversion is clearly present during effort-based decision-making and it can be modulated by altered dopaminergic state. This presents interesting future questions surrounding clinical disorders that have shown a reduced willingness to exert effort such as depression and stroke. For example, it is possible that disorders that have shown a reduced willingness to exert effort in the pursuit of reward could show a normal, or even enhanced, willingness to exert effort in order to avoid punishment.

## Acknowledgments

This work was supported by a European Research Council starter grant to JMG (MotMotLearn: 637488). Participants with Parkinson’s disease were recruited via Parkinson’s UK (www.parkinsons.org.uk).

